# Spire stimulates nucleation by Cappuccino and binds both ends of actin filaments

**DOI:** 10.1101/773812

**Authors:** Alexander O. Bradley, Christina L. Vizcarra, Hannah M. Bailey, Margot E. Quinlan

## Abstract

An actin mesh fills both mouse and fly oocytes. The meshes are built by a conserved mechanism and used to establish polarity. Two actin nucleators, Spire and Cappuccino, collaborate to build actin filaments that connect vesicles and the cortex. Direct interaction between Spire and Cappuccino is required for in vitro synergistic actin assembly; however, we understand little about why the interaction is necessary. To mimic the geometry of Spire and Cappuccino in vivo, we immobilized Spire on beads. We found that increased nucleation is a major part of synergy and that Spire alone binds both barbed- and pointed-ends of actin filaments. We identified Spire’s barbed-end binding domain. Partial rescue of fertility by a loss-of-function mutant indicates that barbed-end binding is not necessary for Spire’s in vivo function, but that it may play a role under normal circumstances. We propose that Spire stimulates nucleation by Cappuccino in a manner similar to the collaboration between APC and mDia1.

**Summary:** Actin nucleators Cappuccino and Spire collaborate to build an actin mesh in oocytes. Data demonstrate that the collaboration leads to synergistic actin nucleation, as opposed to elongation. Further, Spire binds both ends of polar, actin filaments, resolving a long-outstanding question.

## Introduction

Cells contain a variety of actin-based structures that fulfill distinct functional roles. The actin cytoskeleton is malleable due to dynamic structural regulation by a range of distinct actin-binding proteins. The first step to building a structure is generally catalysis of new actin filaments, so-called nucleation. Both a kinetic barrier and certain actin binding proteins, such as profilin, prevent spontaneous nucleation in the cell. Instead, actin nucleators stimulate this process in highly regulated manners. There are three known classes of actin nucleators that function by distinct mechanisms: the Arp2/3 complex, formins, and tandem-WH2 domain nucleators. A developing trend is that none of these proteins work independently. Effector proteins can inhibit or enhance the activity of actin nucleators. Usually, the effector is not a nucleator. In some cases, two independent nucleators also work together. A poorly understood example of such interplay is the collaboration between Cappuccino (Capu, a formin) and Spire (Spir, a tandem-WH2 nucleator).

Both Spir and Capu are required for oogenesis. This discovery was first made in *Drosophila* and subsequently demonstrated in mouse (Manseau and Schupbach, 1989; Leader et al., 2002; Pfender et al., 2011). In both cases, Capu (or Fmn-2, one of two mammalian homologs) and Spir (or mSpire, referring to mammalian homologs, Spire-1 and Spire-2) build a mesh of actin that fills the oocyte (Dahlgaard et al., 2007; Schuh and Ellenberg, 2008; Azoury et al., 2008; Pfender et al., 2011). The discovery that both Spir and Capu are actin nucleators led to the question of why two nucleators would be needed to build one structure (Quinlan et al., 2005). We now know that direct interaction between Spir and Capu is required for their function (Quinlan, 2013). Detailed biochemical analyses and structural biology provide insight into the interaction: the N-terminal Spir-KIND domain binds the C-terminal Capu-tail with ∼100 nM affinity (Fig. 1) (Quinlan et al., 2007; Vizcarra et al., 2011; Pechlivanis et al., 2009; Zeth et al., 2011). However, our understanding of the functional consequences of the interaction remain incomplete. Capu, like all formins, is a dimer. It can bind two copies of Spir and, thereby, accelerate actin assembly by Spir (Quinlan et al., 2007; Vizcarra et al., 2011). In contrast, Spir’s KIND domain inhibits nucleation by Capu and competes with barbed ends for Capu binding, effectively inhibiting Capu’s ability to accelerate actin assembly (Quinlan et al., 2007; Vizcarra et al., 2011). However, when a human Spire-1 construct, containing both the KIND domain and four tandem WH2 domains, is mixed with the C-terminal half of Fmn-2, actin assembly is greatly enhanced (Montaville et al., 2014). So-called ping-ponging – Spir and Fmn-2 alternately binding to the barbed end of filaments – is observed and was proposed to account for synergistic actin assembly (Montaville et al., 2014).

**Fig. 1.**
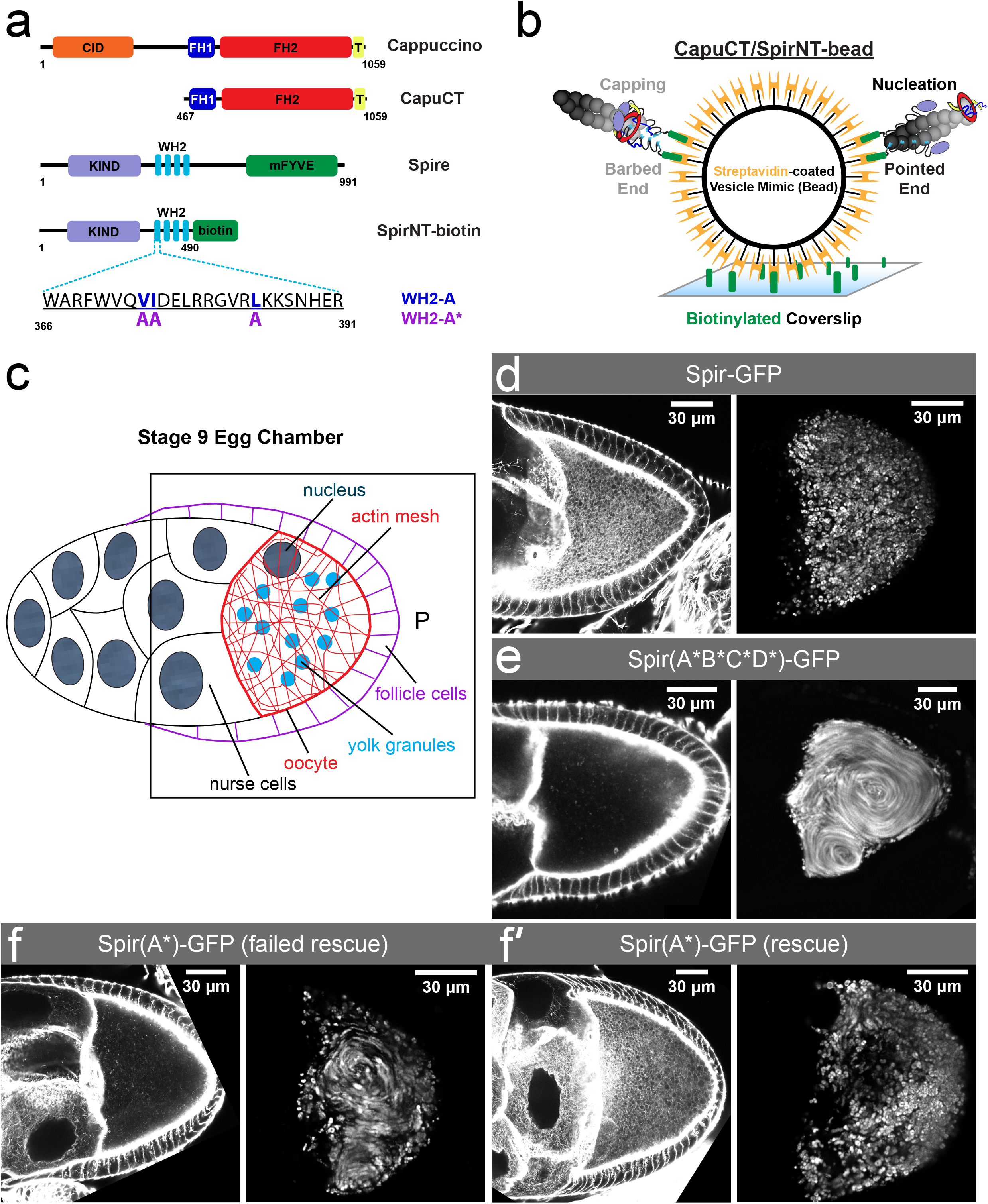
WH2 domains are necessary for oogenesis. *(A)* Domain architecture of *Drosophila melanogaster* Cappuccino (Capu), Spire (Spir), and truncations used. CID, Capu Inhibitory Domain (orange); FH1/FH2, Formin Homology domains (1, dark blue; 2, red); T, Tail (yellow); KIND, Kinase Noncatalytic C-lobe Domain (purple); WH2, Wiskott-Aldrich syndrome Homology domains (light blue); mFYVE, Modified FYVE (zinc finger) domain (green). Spir has four WH2 domains, named WH2-A through -D. The sequence of WH2-A and mutations that disrupt its actin binding are shown. *(B)* Bead experiment methodology. To decorate streptavidin-coated microspheres (gold) with SpirNT, and to bind the microspheres to the coverslip, biotin (green) is conjugated both to coverslips and to SpirNT. Two potential orientations of the actin filament distinguish Spir’s primary activities at each end: capping filaments at their barbed ends (left) and nucleating filaments from their pointed ends (right). (*C*) Cartoon of a stage 9 egg chamber. P, Posterior. (*D – F*) Stage 9 egg chambers dissected from flies with a *spir* null background, expressing Spir-GFP (*D*), Spir(A*B*C*D*)-GFP (*E*), or Spir(A*)-GFP (*F, F’*). Egg chambers were stained with fluorescently-labeled phalloidin to detect the presence or absence of mesh (left images) and standard deviation projections of autofluorescent yolk granule positions over 2 minutes (right images) show the extent of ooplasmic streaming. Expression of Spir(A*)-GFP results in a mixture of failed (*F*) and successful rescues (*F’*).

Spir is enriched at the cortex of the *Drosophila* oocyte, localizes to Rab11-positive vesicles in mouse oocytes, and the C-terminal mFYVE domain of mSpire-1 binds phospholipids (Quinlan et al., 2007; Quinlan, 2013; Pfender et al., 2011; Tittel et al., 2015). While Fmn-2 is also observed on Rab11-positive vesicles in mouse ooctyes, GFP-Capu appears diffuse throughout the *Drosophila* oocyte (Schuh, 2011; Dahlgaard et al., 2007; Quinlan, 2013). In the mouse oocyte, Rab11-positive vesicles that contain mSpire, Fmn-2, and Myosin V are at nodes of the actin mesh that fills the oocyte (Schuh, 2011). Mesh dynamics contribute to nucleus positioning and long distance transport of the Rab11-positive vesicles toward the cell cortex (Holubcová et al., 2013; Almonacid et al., 2015; Ahmed et al., 2018). A model to account for the long distance transport includes mSpire/Fmn-2-nucleated actin filaments growing with their barbed ends remaining near the vesicles, creating tracks that myosin V on neighboring vesicles can walk along, pulling vesicles together (Schuh, 2011). When ping-ponging was observed, Montaville et al. (Montaville et al., 2014) expanded on this model, proposing that barbed ends of filaments and/or pre-nuclei are recruited by vesicle-bound mSpire and handed off to Fmn-2 to stimulate elongation.

In order to learn more about how Spir and Capu interact to build actin filaments and structures, we combined biochemistry and fly genetics. We developed a bead-based assay, in which we attached the N-terminal half of Spir (SpirNT) to beads and added free C-terminal half of Capu (CapuCT), based on reported localization data (see Fig. 1A for construct definitions). Through this work we found that beads decorated with SpirNT and CapuCT (CapuCT/SpirNT-beads) nucleate filaments that grow with their barbed ends away from beads, protected by CapuCT, opposite to the polarity proposed earlier. We found that SpirNT alone on beads (SpirNT-beads) nucleates filaments oriented the same way. We also found that SpirNT-beads retain the pointed ends of filaments with a dwell time of ∼100 seconds while neighboring SpirNT-beads capture the barbed ends of filaments for several hundreds of seconds. These data resolve an outstanding question in the literature about Spir’s association with filament ends – it binds both ends. We also identified the domain necessary for Spir’s high-affinity barbed-end binding. Surprisingly, barbed-end binding is not necessary for oogenesis and loss of barbed-end binding increases Spir/Capu synergy *in vitro*. These genetic and biochemical data indicate that ping-ponging at the barbed-end is neither the dominant source of synergy nor necessary *in vivo*. Instead, we propose that Spir and Capu collaborate by a mechanism in which Spir promotes Capu’s nucleation activity through its independent nucleation activity, similar to APC/mDia1 synergy (Okada et al., 2010; Breitsprecher et al., 2012).

## Results

### WH2 domains are necessary for Drosophila oogenesis

The Spir-KIND domain inhibits CapuCT in pyrene-actin assays (Quinlan et al., 2007; Vizcarra et al., 2011). However, genetics indicate that Spir and Capu both play positive roles in oocyte mesh formation (Manseau and Schupbach, 1989; Dahlgaard et al., 2007; Quinlan, 2013). Combined, these data suggest that Spir’s WH2 domains contribute to function in vivo. To formally test whether actin binding is required, we used full-length Spir (CG10076, splice variant PB) with a C-terminal mEGFP tag in a genetic rescue assay. In each of Spir’s four WH2 domains, we mutated three conserved, hydrophobic residues which contact actin (Fig. 1A) (Kelly et al., 2006; Chereau et al., 2005). These mutations were previously shown to dramatically decrease, if not abolish, actin monomer binding (Quinlan et al., 2005). In vitro, SpirNT with all of these mutations (SpirNT(A*B*C*D*)) does not accelerate actin assembly (Fig. S1A) (Quinlan et al., 2005). We previously demonstrated that expression of wild-type Spir-GFP, driven by germline specific nanos-Gal4-vp16, is sufficient to rescue fertility in flies that lack endogenous Spir (Table 1) (Quinlan, 2013). In contrast, expression of Spir(A*B*C*D*)-GFP failed to rescue fertility (hatch rate < 2%, Table 1). In oocytes expressing only Spir(A*B*C*D*)-GFP, the actin mesh is absent and streaming is premature, consistent with loss of actin assembly activity and loss of fertility (Fig. 1C-E). Thus, we conclude that actin binding by at least one of Spir’s WH2 domains is necessary for actin assembly and oogenesis.

**Table 1.** Fertility Assays.

| Transgene (*spir*<sup>1</sup>/*Df*(2L)*Exel*<sup>6046</sup>)<sup>a</sup> | % Hatched<sup>b</sup> | Number counted<sup>b</sup> |
| --- | --- | --- |
| *Spir*-GFP<sup>c</sup> | 59 | 606 |
| *Spir*(A*B*C*D*)-GFP | <2 | 431 |
| *Spir*(A*)-GFP | 40 | 619 |

<sup>a</sup>Genetic background is in parentheses.
<sup>b</sup>% hatched is reported as the average of at least three independent trials. Number counted is the sum of eggs counted from all trials.
<sup>c</sup>(Quinlan, 2013)
% hatched is measured 24 hours after egg lay. ~95% w<sup>1118</sup> hatch; 0% *spir*<sup>1</sup>/*Df*(2L)*Exel*<sup>6046</sup> hatch.

### Barbed ends project away from CapuCT/SpirNT-beads

In order to learn more about how Spir and Capu interact to build actin filaments and structures, we developed a bead-based assay. Spir is enriched at the cortex of the *Drosophila* oocyte, localizes to Rab11-positive vesicles in mouse oocytes, and the C-terminal mSpire-mFYVE domain binds phospholipids (Quinlan et al., 2007; Quinlan, 2013; Pfender et al., 2011; Tittel et al., 2015). While Fmn2 is also observed on vesicles in the mouse oocyte, GFP-Capu appears diffuse throughout the *Drosophila* oocyte (Schuh, 2011; Quinlan, 2013). Based on these data, we immobilized SpirNT on beads and added CapuCT in solution. To do so, we biotinylated a C-terminal Avitag on SpirNT and mixed it with CapuCT and streptavidin-coated beads (Fig. 1B). We introduced the decorated beads to a sparsely biotinylated flow chamber and imaged them by total internal reflection fluorescence (TIRF) microscopy. When we added actin (20% OG-actin) and profilin to the flow chamber, extensive and rapid polymerization followed (Fig. 2A**, Video 1**).

**Fig. 2.**
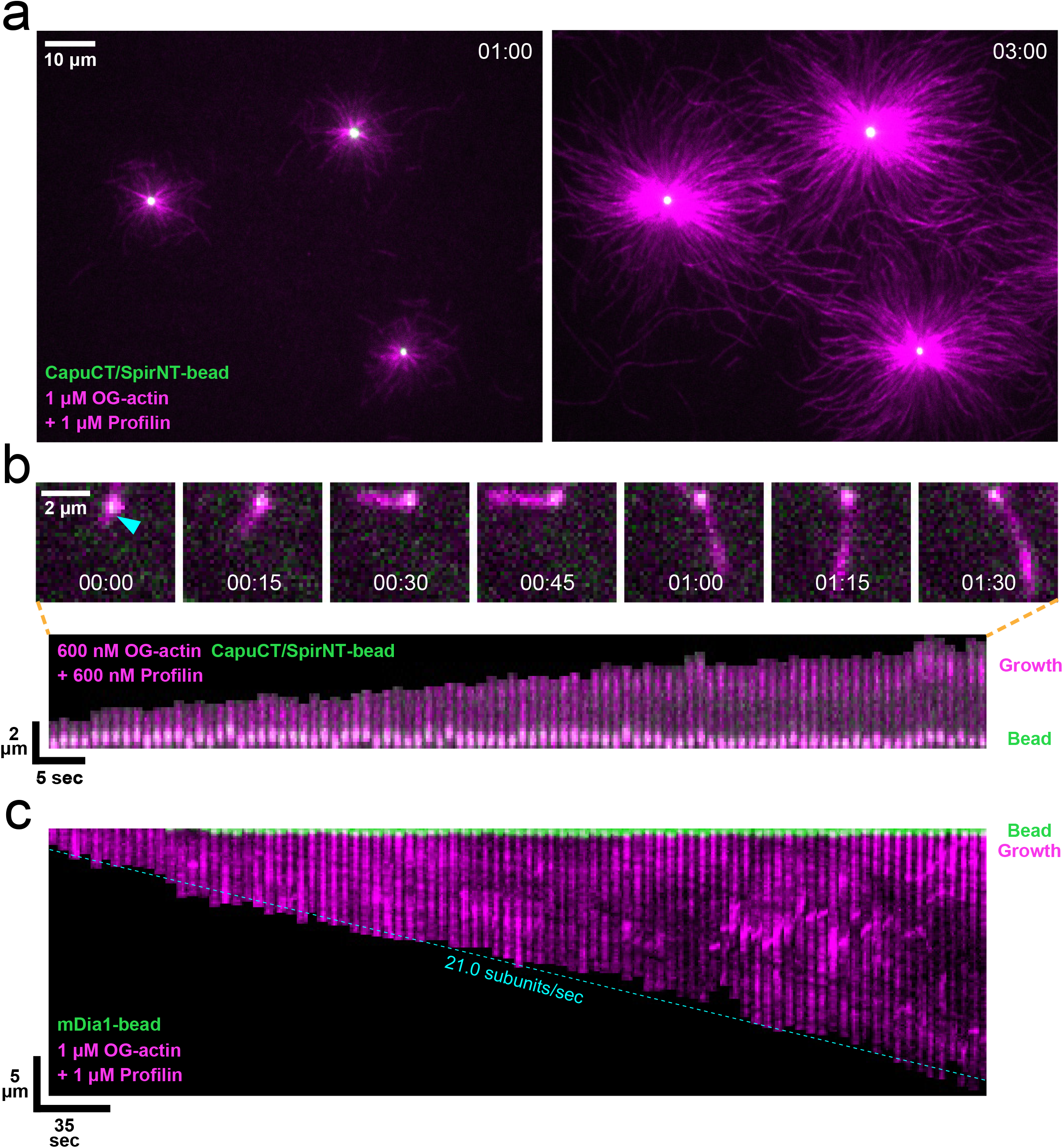
Barbed ends project away from CapuCT/SpirNT-beads. *(A)* Actin assembles off of CapuCT/SpirNT-beads. See also, Video 1*. (B)* A single filament is nucleated and retained by a bead. Kymograph (bottom) shows bleaching of filament regions closest to the bead and new, labeled actin added away from the bead, indicating growth away from the bead. See also, Video 3*. (C)* A single filament is nucleated and elongated by a bead coated with mDia1. Filament growth is accelerated by the processive formin and profilin. Fiducial marks (e.g. dark, diagonal lines) in the kymograph are displaced from the bead at the rate of elongation, indicating growth at the bead. See also, Video 4.

Two groups proposed that Spir/Capu-coated vesicles have filaments with barbed ends apposed to the vesicle and pointed ends growing away (Fig. 1B) (Montaville et al., 2014; Schuh, 2011). We tested this model by examining the orientation of filaments growing off of CapuCT/SpirNT-beads. By lowering the concentration of actin (600 nM), we decreased nucleation such that we could track individual filaments. Under these conditions, we observed filaments growing from but “retained” at the beads (**Videos 2 and 3**). We carefully analyzed 9 of 32 individual filaments observed. In every case, fiducial marks which do not move with respect to the bead, and increased fluorescence intensity at filament ends (i.e. ends brighter due to addition of unbleached monomers (Kovar and Pollard, 2004)), away from the bead, indicate that barbed ends grow away from the bead surface (Fig. 2B**, Video 3**). We expected to see the opposite case with Capu directly conjugated to the beads. For unknown reasons, Capu was not functional when attached to beads through a number of different linkers. Instead, we conjugated the formin mDia1 to beads. In this case, fiducial marks were displaced at the rate of filament elongation and the addition of unbleached monomers was no longer observed at the filament end away from the bead (Fig. 2C**, Video 4**), consistent with barbed-end growth at the bead surface. Together these data demonstrate that CapuCT/SpirNT-beads nucleate filaments with barbed ends growing away from the bead and suggest that CapuCT separates from SpirNT to elongate the filament.

### Spir beads retain the pointed ends of nucleated actin filaments

A dense collection of filaments emanated radially from CapuCT/SpirNT-beads. Because we concluded that CapuCT-mediated elongation proceeded away from the beads, the density of actin near the bead surface suggested that the pointed ends of filaments could be retained by Spir. We repeated the experiments with SpirNT-beads, without added CapuCT or profilin, and observed similar patterns (Fig. 3A**, Video 5**). We found that nucleation by SpirNT-beads was suppressed enough to observe individual filaments when profilin was added back (Fig. 3B). We again observed filaments growing from and retained at the bead surface (Fig. 3C **and Video 6**). The distribution of filament dwell times was well fit by a single exponential with an off rate of 0.007 ± 0.003 s^-1^ (Fig. 3D). We note that retention of filaments by SpirNT-beads occurred independent of profilin (Fig. S1G **and Video 7**). Furthermore, neither filament end was observed to interact with beads coated with Spir-KIND and CapuCT (Fig. S1F), indicating that retention is specifically mediated by Spir-WH2 domains.

**Fig. 3.**
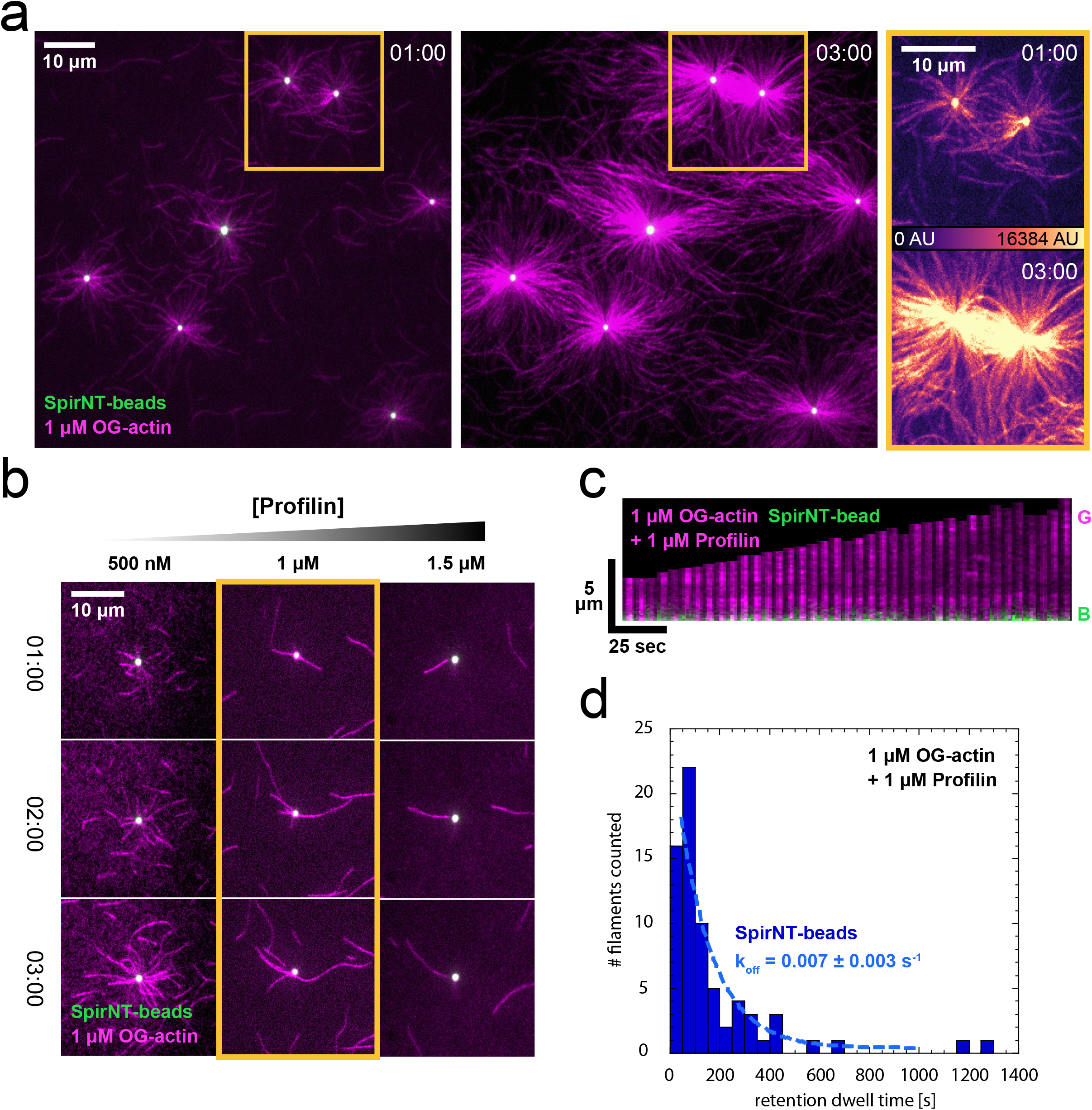
SpirNT-beads retain the pointed ends of nucleated actin filaments. *(A)* SpirNT-beads, in the absence of Capu and profilin, also potently polymerize actin. The highlighted region (gold) is magnified and pseudocolored (right) to emphasize the enrichment of actin filaments between beads. See also, *Video 5. (B)* Nucleation by SpirNT-beads is suppressed by profilin. When added at 1 μM (gold box), profilin permits the observation of several, single filaments per bead. *(C)* SpirNT-beads also retain nucleated filaments. G, Growth; B, Bead. Fiducial marks (dark, horizontal lines) in the kymograph are not displaced as the filament elongates, indicating growth away from the bead. See also, *Video 6. (D)* Dwell times of filament pointed ends on beads. Nucleated and retained filaments were tracked from SpirNT-beads in the presence of 1 μM actin and 1 μM profilin. All bead-associated filaments were counted unless obviously captured from solution or > 1 μm in length when first visible. The data are well fit by a single exponential (k_off_ = 0.007 ± 0.003 s^-1^, n = 70 filaments; 4 independent experiments).

We did not assume that filaments were oriented with their barbed ends out when nucleated by SpirNT-beads because Spir has been reported to bind both the barbed and pointed ends of filaments (Quinlan et al., 2005; Bosch et al., 2007; Ito et al., 2011). mSpire1 caps barbed ends with high affinity and SpirNT binds pointed ends weakly (nM vs μM K_d_s) (Bosch et al., 2007; Montaville et al., 2014; Quinlan et al., 2005). We closely analyzed 10 of the 70 filaments included in the retention data set. All ten filaments, growing from SpirNT-beads, displayed fiducial marks which did not move with respect to the bead and typically had brighter filament ends away from the beads consistent with barbed ends being oriented away from the beads (Fig. 3C). Thus, we conclude that barbed ends grow away from SpirNT-beads and that SpirNT is sufficient to retain the pointed ends of actin filaments for >100 seconds.

### SpirNT-beads capture and cap the barbed ends of actin filaments

When we examined the dense actin networks emanating from beads, we also observed apparent connections between neighboring beads (Fig. 3A). This pattern suggested to us that the barbed ends of actin filaments could also be associated with the beads. Under conditions where individual filaments could be tracked, we observed the “capture” of filaments (Fig. 4). The association of captured filaments with Spir beads often outlasted the single-filament imaging window (10+ minutes), precluding measurement of an off rate and suggesting that the interaction was distinct from retention.

**Fig. 4.**
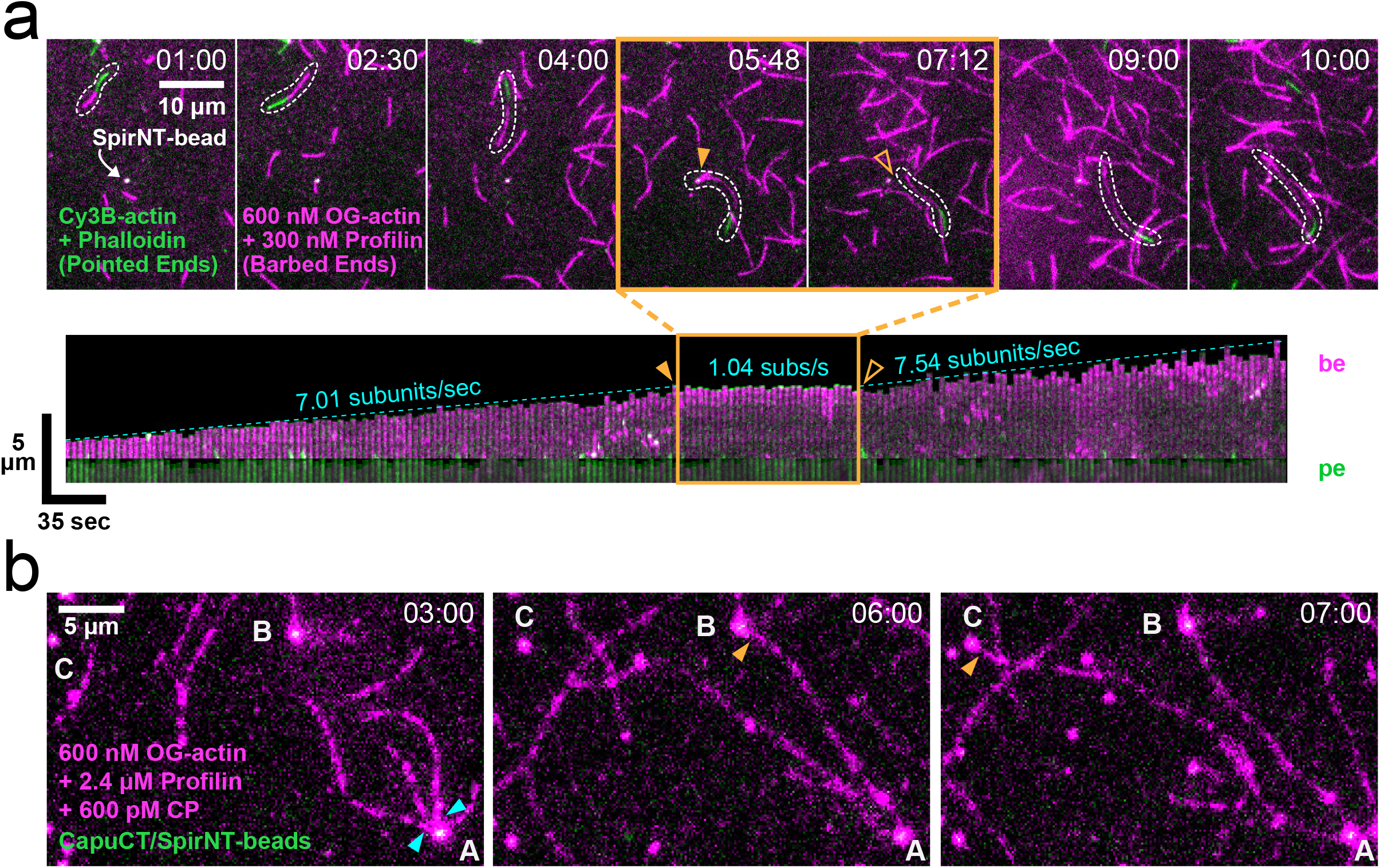
SpirNT-beads capture and cap the barbed ends of actin filaments. *(A)* Nucleated filaments (green) grow only at their barbed ends (magenta). A free filament (dashed outline) diffuses and grows until it is captured by a SpirNT-bead (solid triangle). The filament resumes growth once released by the bead (open triangle). The kymograph shows that the growing, barbed end (magenta) is captured/capped by Spir and resumes growth after release. See also, *Video 8. (B)* Two filaments (cyan triangles) nucleated by a SpirNT/CapuCT-bead (“A”) are retained for over 5 minutes. Their barbed ends are captured (gold triangles) by other CapuCT/SpirNT-beads (“B” and “C”). Accelerated growth of these filaments (∼20 subunits/sec) in the presence of profilin and capping protein indicates that CapuCT is elongating and protecting their barbed ends. Filaments did not measurably grow, following capture, suggesting that CapuCT is displaced and that the barbed end is bound by SpirNT upon capture. See also, *Video 9*.

In addition to seeing filaments between beads, we often observed capture of free filaments. We probed the orientation of captured filaments using two colors of actin. We initiated elongation in the presence of Cy3B-actin and stabilized these filaments with phalloidin after several minutes. Adding a limiting pool of OG-actin (600 nM) plus profilin to favor barbed end growth, we observed that – whether the filament had both ends free (as shown) or one end retained by a bead – only the barbed ends of filaments were captured by beads (Fig. 4A**, Video 8**). Filaments captured by beads did not grow measurably (Fig. 4A). Occasionally, we observed the release of a barbed end, following capture. In these cases, the filament resumed elongation at the same rate as before capture (Fig. 4A). As noted above, barbed ends do not interact with CapuCT/Spir-KIND-beads (Fig. S1F). These data indicate that the barbed ends are capped by Spir-WH2 domains, as opposed to being non-specifically stuck to the beads or KIND domain.

To further test the phenomenon of filament capture, we added capping protein and CapuCT to the experiment. Under these conditions, the sustained and accelerated growth of retained filaments confirmed that CapuCT separates from SpirNT to elongate filament barbed ends (Fig. 4B). These filaments were still captured by beads, suggesting that SpirNT displaces CapuCT from the barbed end (Fig. 4B**, Video 9**). Together these data demonstrate that – with or without CapuCT present – Spir-beads can nucleate and retain the pointed ends of actin filaments, while neighboring beads can capture and cap their barbed ends.

### Spir’s WH2-A binds filament barbed ends and reduces actin nucleation

Each of Spir’s four WH2 domains contributes differently to nucleation and they bind actin monomers with a range of affinities (from ∼100 nM – 1.1 μM) (Quinlan et al., 2005; Rasson et al., 2015). We reasoned that the most N-terminal WH2 domain, WH2-A, binds barbed ends of filaments, based on our earlier observation that isolated WH2-A is the only Spir-WH2 domain that slows filament growth (Rasson et al., 2015). To test the contribution of WH2-A to barbed end binding, we first established that wild-type SpirNT caps barbed ends in a seeded elongation assay, as has been shown for mSpire-1 (Bosch et al., 2007). In the presence of actin seeds, monomers, and profilin, SpirNT potently inhibits elongation (Fig. 5A). The dose dependence of inhibition indicates an apparent affinity of SpirNT for barbed ends of 20 ± 3 nM, similar to that reported for mSpire-1 (Fig. 5B). We then mutated three conserved hydrophobic residues in WH2-A of SpirNT (SpirNT(A*); Fig. 1A) to remove actin monomer binding by this domain alone. When added to the seeded elongation assay, SpirNT(A*) was essentially unable to inhibit elongation (Fig. 5A,B). Thus, a functional WH2-A domain is necessary for high affinity, barbed-end capping.

**Fig. 5.**
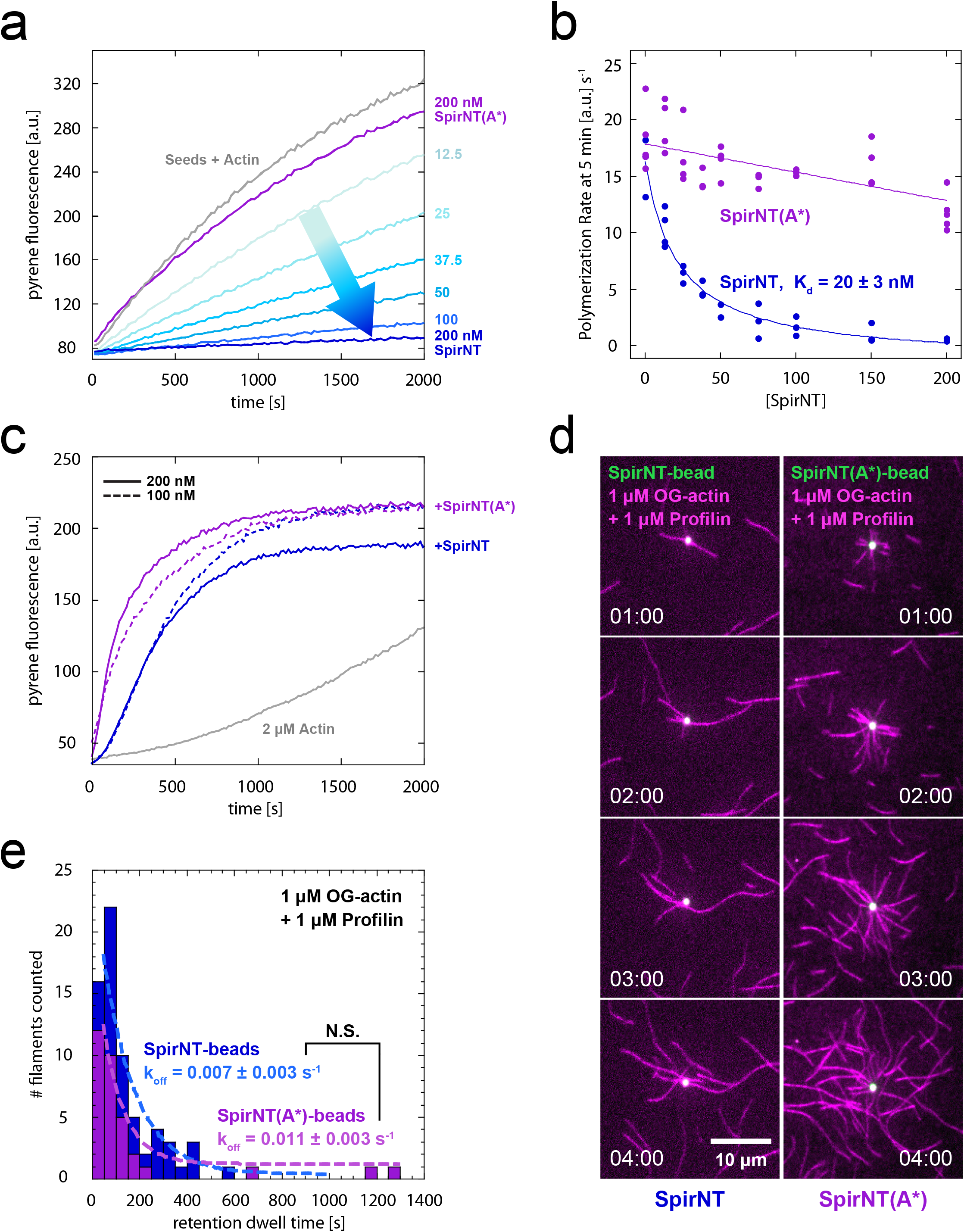
Spir’s WH2-A binds filament barbed ends and reduces actin nucleation. *(A)* Seeded elongation is inhibited by the addition of 12.5 – 200 nM SpirNT (shades of blue). The addition of 200 nM SpirNT(A*) has no effect (purple). *(B)* Dose dependent elongation rates are plotted for three experiments. The data are fit by a quadratic equation, indicating that SpirNT binds the barbed end with a K_d_ of 20 ± 3 nM. The line is a fit to all data points and the K_d_ is the average of three independent experiments. Inhibition by SpirNT(A*) is negligible. *(C)* SpirNT(A*) assembles actin more potently than wild type SpirNT in a pyrene-actin assembly assay. Plateaus that are independent of Spire concentration suggest that SpirNT(A*) does not sequester actin like SpirNT (compare solid lines). *(D)* SpirNT(A*)-beads nucleate more potently than wild type. See also, *Video 10. (E)* Quantification of filament pointed end dwell times on beads (k_off_ = 0.011 ± 0.003 s^-1^,n = 33 filaments; 3 independent experiments). The off rate is not statistically different from wild type (data from Fig. 3D).

In standard pyrene-actin assays, we found that SpirNT(A*) assembles actin more potently than wild type SpirNT (Fig. 5C **and** SC). Of note, the plateau of pyrene traces does not decrease at high concentrations as is seen for wild type SpirNT, suggesting that this mutant is not a potent monomer sequesterer (Fig. 5C) (Quinlan et al., 2005). We next attached biotinylated SpirNT(A*) to beads and added actin and profilin. Consistent with increased activity in pyrene assays, we observed faster accumulation of filaments from SpirNT(A*)-beads compared to wild type SpirNT-beads (Fig. 5D**, Video 10**). To compare nucleation rates on beads, we measured the integrated intensity of actin within a band 1.6 μm away from the beads. At this proximity, we are minimally sensitive to filament elongation and interpret the intensity as proportional to the number of filaments; that is, nucleation. We measured time courses and calculated the rates of increase in OG-actin signal. The difference in rates indicates that nucleation by SpirNT(A*) is ∼3x stronger than wild type SpirNT in the presence of profilin (Fig. 6D,E). We also measured the dwell times of filaments retained by SpirNT(A*)-beads. The off rate is not significantly different than that of wild type (k_off(**WT**)_ = 0.007 ± 0.003 s^-1^, k_off(**A\***)_ = 0.011 ± 0.003 s^-1^, Student’s t-test, p = 0.98, Fig. 5E), indicating that WH2-A does not play an important role in filament retention. Notably, we rarely observed apparent capture events by SpirNT(A*)-beads. In several cases, filaments appeared to be close enough to a neighboring bead for capture for tens of seconds, but sustained association of the barbed end with beads was rare. Taken together, these data are consistent with the requirement of functional WH2-A for high affinity barbed end binding and demonstrate that this domain is not critical for pointed end binding.

**Fig. 6.**
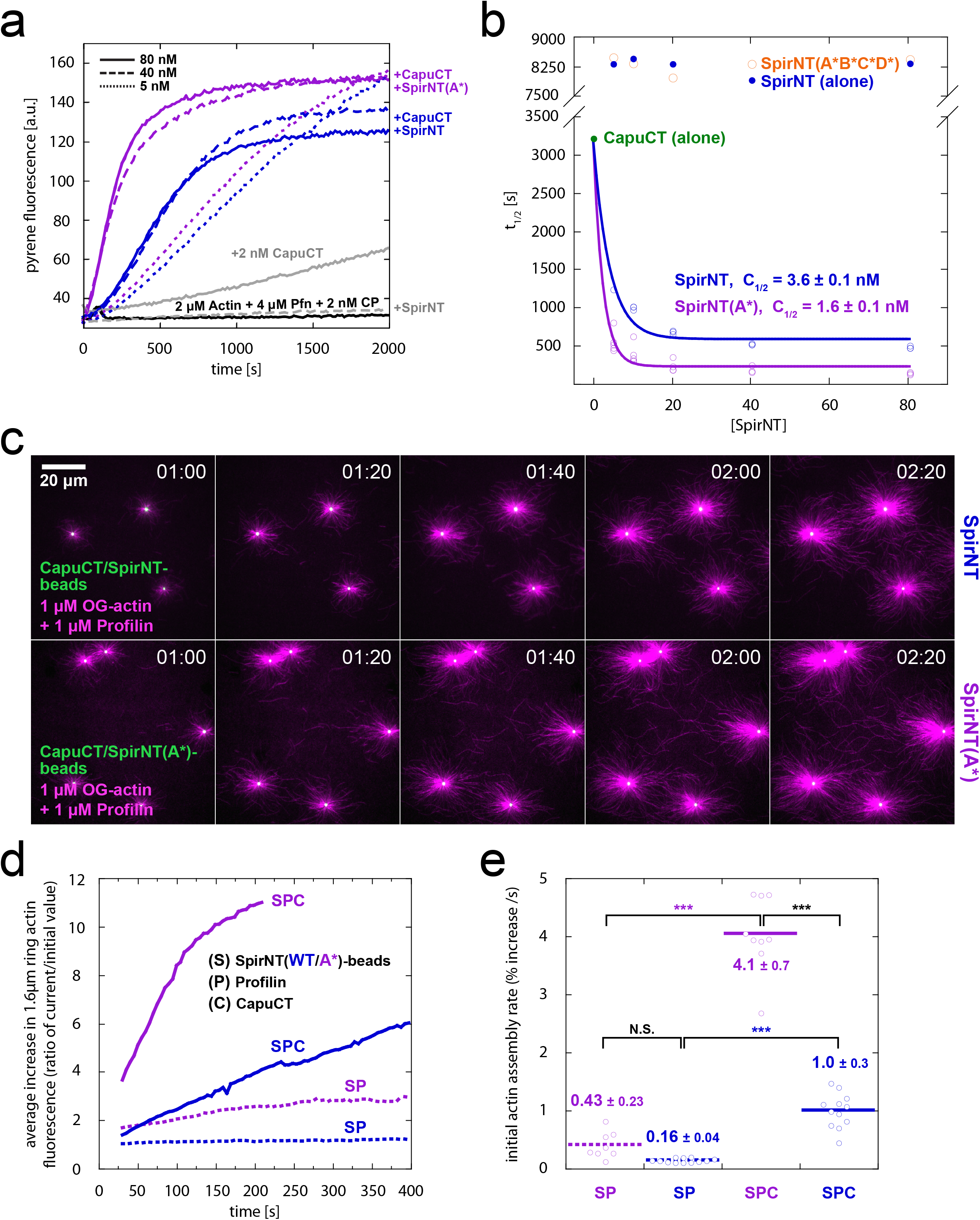
Synergy between Spir and Capu does not require barbed end binding. *(A)* Actin assembly assays in the presence of profilin and capping protein. CapuCT alone and SpirNT alone (solid and dashed gray lines) are weak under these conditions, but synergize in assembly when combined (blue). SpirNT(A*) synergizes more potently with CapuCT (magenta). *(B)* Dose dependence of t_1/2_. Solid circles are SpirNT or CapuCT alone (blue and green, respectively; one representative experiment shown for each). Open circles indicate the addition of both proteins. SpirNT, blue; SpirNT(A*), magenta (n = 3 independent experiments, each); SpirNT(A*B*C*D*), orange (one representative experiment shown at each concentration). Raw data are shown in Fig. S1B,D. *(C)* Nucleation is stronger on CapuCT/SpirNT(A*)-beads than CapuCT/SpirNT-beads. *(D)* Quantification of experiments like those shown in *(C)* and Fig. 5D. Fluorescence around beads is expressed as an average percentage of initial values over time (n ≥ 4 beads, n ≥ 2 independent experiments per condition). See methods for description of data transformation. Data plotted were obtained using 1 μm diameter beads, 1 μM actin, and 1 μM profilin, with or without CapuCT (solid or dashed lines, respectively). S, SpirNT-beads; P, profilin; C, CapuCT. SpirNT-bead data are plotted in blue and SpirNT(A*)-bead data, in magenta. (*E*) Rates of fluorescence increase, from (*D*). Mean ± SEM are indicated for each condition. The initial 50 s of images (t = ∼30–80 s) were used to determine rates. A Student’s t-test was performed on each pair of conditions indicated (black brackets). ***, p < 0.0001. The difference in activity by SpirNT(A*/WT)-beads (without Capu) was not statistically significant (unequal variance; p = 0.13).

### Synergy between Spir and Capu does not require barbed end binding

Previously, synergistic actin assembly by mSpire-1 and Fmn-2 was described: the rate of actin assembly by mSpire-1+Fmn-2 is greater than the sum of assembly rates of the individual nucleators (Montaville et al., 2014). We confirmed that SpirNT and CapuCT exhibit a similar synergistic effect in pyrene-actin assembly assays (Fig. 6A **and** S1D). In the presence of profilin, the assembly rate of SpirNT+CapuCT (based on the t_1/2_ at ≥ 40 nM SpirNT) was 6x and15x faster than CapuCT or SpirNT, respectively (Fig. 6B). As seen for the mammalian paralogs, direct interaction is required for synergy: SpirNT(Y232K), a mutant that does not bind Capu, does not synergize with CapuCT (Vizcarra et al., 2011; Montaville et al., 2014) (Fig. S1E). Likewise, Spir’s actin binding activity is required, as demonstrated by the absence of synergy when SpirNT(A*B*C*D*) is added to CapuCT (Fig 6B **and** S1B).

Montaville et al. (Montaville et al., 2014) concluded that actin assembly was enhanced by mSpire-1 and Fmn-2 alternately binding the barbed end, which they dubbed the “ping-pong” mechanism. Given that SpirNT(A*) does not bind to or capture barbed ends, we assume that ping-ponging is not possible with this mutant. However, when we tested SpirNT(A*) in the pyrene synergy assay, we found ∼3x enhanced synergy (based on the t_1/2_ at ≥ 40 nM SpirNT), as opposed to loss of activity (Fig. 6A,B **and** S1D). The dose dependence of synergy is indistinguishable for SpirNT and SpirNT(A*) (Fig. 6B). We speculate that dose dependence is a function of Spir-KIND/Capu-tail binding, which should not be affected in SpirNT(A*). These data lead us to propose that nucleation – not just elongation – is enhanced when SpirNT and CapuCT interact.

To test more directly whether nucleation is increased by SpirNT and CapuCT collaborating, we returned to the bead assay. When CapuCT/SpirNT-beads are mixed with actin and profilin, we observe potent nucleation (Fig. 6C). Consistent with nucleation being a significant element of Spir/Capu synergy, the rate of filament formation is more than 6x greater than nucleation in the absence of Capu (Figs. 6C,D,E). Nucleation of profilin-actin by CapuCT/SpirNT(A*)-beads was even stronger than wild type (4x) and the synergy more pronounced (9.5x vs SpirNT(A*)-beads in the absence of Capu) with the actin signal saturating at the measured 1.6 μm radius within ∼2 minutes (Fig. 6C,D). Thus, nucleation is potently enhanced when SpirNT and CapuCT are combined and loss of barbed end binding by SpirNT leads to even stronger nucleation in the presence and absence of CapuCT. These data support our conclusion that enhanced nucleation is a major source of synergy.

### Barbed end binding is not necessary for oogenesis

Finally, we asked whether barbed-end binding is necessary for *Drosophila* oogenesis. To do so, we tested Spir(A*)-GFP, using the rescue strategy described above. We found that, when driven by nanos-Gal4-vp16, fertility was partially rescued (40%, Table 1). The fertility rescue is about two thirds as effective for Spir(A*)-GFP when compared to the wild type transgene. Consistently, the mesh is present and streaming is properly regulated in about half of the oocytes (Fig. 1F,F’) Thus, barbed-end binding is not necessary for mesh formation or oogenesis, though the fertility decrease suggests that it may contribute under normal conditions.

## Discussion

### Spir binds both ends of actin filaments

The bead assay facilitated observation of several steps in actin assembly by Spir and Capu. We found that filaments nucleated on CapuCT/SpirNT-beads grow with their barbed ends away from the bead with enhanced rates, accelerated by CapuCT. Thus, SpirNT and CapuCT separate after nucleation, consistent with genetics results (Quinlan, 2013). We also found that CapuCT-bound barbed ends would subsequently stop growing if they encountered CapuCT/SpirNT-beads, suggesting that the barbed-end was passed from CapuCT to SpirNT. To our surprise, we found that SpirNT-beads were sufficient to bind both ends of actin filaments. There are conflicting reports regarding filament-end binding for two classes of tandem-WH2 nucleators: Spir and VopF/L (Liverman et al., 2007; Yu et al., 2011; Pernier et al., 2013, 2016; Burke et al., 2017). The N-termini of WH2 domains bind actin monomers between subdomains 1 and 3 of actin – the surface exposed at the barbed ends of filaments (Chereau et al., 2005; Hertzog et al., 2004). It was, therefore, reasonable to expect tandem-WH2 nucleators to associate with filament barbed ends and surprising when both VopL and Spir were reported to bind filament pointed ends (Quinlan et al., 2005; Namgoong et al., 2011; Yu et al., 2011). Spir nucleates filaments with free barbed ends and inhibits depolymerization of gelsolin-capped filaments, albeit weakly (Quinlan et al., 2005). Spir also caps barbed ends of growing filaments but with nanomolar affinity (Bosch et al., 2007; Montaville et al., 2014). Here we present evidence that these apparently conflicting data are both correct. This is not unprecedented. Namgoong et al. (Namgoong et al., 2011) and Burke et al. (Burke et al., 2017) reported that VopL/F can interact with both ends of actin filaments, depending on the conditions.

Earlier, we did not observe barbed-end binding by Spir when assayed by inhibition of depolymerization (Quinlan et al., 2005). We now report high affinity barbed-end binding in inhibition of elongation assays. Possibly, Spir only binds barbed ends in the presence of actin monomers as was described for N-Wasp (Co et al., 2007). This contrasts with VopL/F that primarily bind barbed ends of preformed filaments in the absence of free monomer (Burke et al., 2017). The first of Spir’s tandem WH2 domains, WH2-A, is necessary to cap barbed ends. It is curious that removing function of this WH2 domain increases the activity of SpirNT. In earlier work, we did not observe a significant difference in actin assembly, with or without WH2-A (Quinlan et al., 2005). The original data were acquired with higher concentrations of both SpirNT and actin, which may have masked the difference. In addition, the data were acquired with a slightly longer construct: 1-520 vs 1-490. One possible explanation for enhanced actin nucleation by SpirNT(A*) is that loss of capping leads to fewer so-called SA_4_ complexes (Spir bound to four actin monomers). We and others observed SA_4_ complexes when Spir is mixed with actin under polymerizing conditions (Quinlan et al., 2005; Bosch et al., 2007). We originally interpreted these structures as pre-nuclei. In contrast, Bosch (Bosch et al., 2007) proposed that they are stable structures that sequester actin monomers. Whether there are two paths (i.e. nucleation vs. SA_4_ complex), or one (nucleation, with formation of the nucleus from the SA_4_ complex being a rate limiting step), removing capping would likely destabilize the SA_4_ structure and could favor nucleation.

How do tandem-WH2-domain nucleators bind filament pointed ends? Single molecule observations showed that VopL/F remains associated with the pointed ends of filaments it nucleates for ∼100 s (Burke et al., 2017). In the absence of actin monomers, VopL/F binds pointed ends for shorter times (∼25 s). Higher affinity binding is likely due to the contribution of the VopL/F C-terminal domain, that dimerizes and binds the pointed end. In the case of Spir, weak inhibition of depolymerization may be mediated by a linker or a domain straddling the WH2 domains. Side binding by the C-terminal portion of tandem WH2 domains may enhance relatively weak pointed end binding of both Spir and VopL/F. Side binding by Spir is consistent with the fact that it can sever filaments (Bosch et al., 2007; Chen et al., 2012). We speculate that the ability to retain the pointed ends of filaments nucleated by Spir on beads reflects enhanced binding due to clustering on beads (Fig. 7A). Multiple cases of emergent behavior of clustered WH2 domains have been reported. Spir nucleates more potently when clustered on gold particles (Ito et al., 2011). VopL retains the pointed end of ∼10% of nucleated filaments for several minutes (compared to seconds for ∼80% of filaments) and accelerates barbed end elongation of ∼8% of filaments when clustered on Qdots (Namgoong et al., 2011); and Ena/VASP are potent elongation factors when clustered on beads (Breitsprecher et al., 2008; Winkelman et al., 2014). In sum, WH2 domains can bind both ends of actin filaments but they are likely to be part of a larger context which dictates their activity (Dominguez, 2016).

**Fig. 7.**
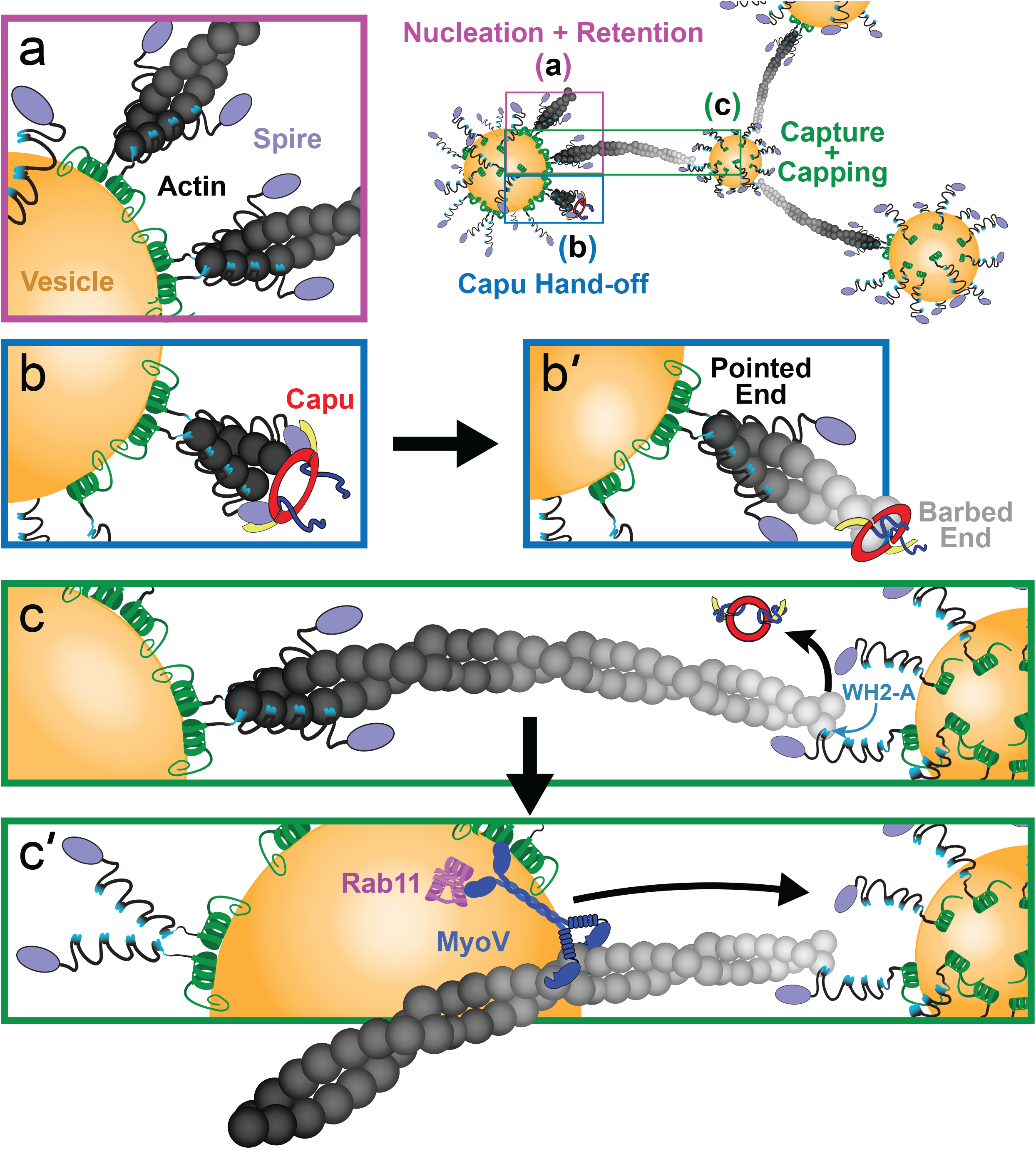
Model of Spir—Capu synergy. *(A)* Spir is directed to vesicles through its phospholipid-binding mFYVE domain (green helix). The clustering of Spir on vesicles may enhance its affinity for filament pointed-ends. Each of Spir’s tandem, WH2 domains (light blue) can bind to an actin subunit (dark gray spheres). *(B)* Spir and Capu interact through their respective KIND (purple) and tail (yellow) domains. *(B’)* Spir retains a filament’s pointed end (dark gray), while Capu separates from Spir to elongate its barbed end (light gray). *(C)* A filament’s barbed end is captured and capped by Spir’s WH2-A domain (light blue), displacing Capu. Spir’s bipolar filament binding may tether vesicles, as shown. *(C’)* Associated with Rab11 (pink) and Spir on vesicles, MyoV (dark blue) may tow a nucleating vesicle (left) toward another, receiving (filament capturing) vesicle (right).

### How do Spir and Capu synergize?

Synergy of actin assembly can be explained by enhanced nucleation and/or elongation. In the cases of Bud6/Bni1 and APC/mDia1, nucleation is enhanced (Graziano et al., 2011; Breitsprecher et al., 2012). We propose that increased nucleation is also the dominant source of synergy for Spir/Capu. The ping-pong model, initially proposed, suggests that Spir and Capu (Fmn2) enhance elongation by dynamic exchange at the barbed ends of filaments. A key element of the ping-pong model is that Fmn2 binds barbed ends with a slow on-rate. Thus, by recruiting Fmn2 to barbed ends, Spir increases the fraction of time that elongation is enhanced by the formin. We found that synergistic assembly of profilin-actin is actually improved in the absence of Spir binding to barbed-ends (CapuCT+SpirNT(A*)). We also directly observed increased nucleation in bead assays: CapuCT/SpirNT-beads nucleate 6x and 10x more potently than SpirNT-beads, with or without a functional WH2-A, respectively (Figs. 5D, 6C-E). A full comparison (i.e. nucleation by SpirNT-beads and CapuCT-beads, versus CapuCT/SpirNT-beads) was precluded by the inactivity of CapuCT-beads, regardless of the coupling method. Finally, we see a 4–fold increase in nucleation by CapuCT/SpirNT(A*)-beads compared to wild type CapuCT/SpirNT-beads, similar to the ∼3x decrease in t_1/2_ in bulk assays (Fig. 6B). Thus, we conclude that nucleation is the major source of enhanced actin assembly mediated by Spir/Capu synergy.

Ping-ponging may still take place and contribute to mesh assembly. We observe transfer of Capu-bound barbed ends to Spir (Fig. 4B, **Video 9**), consistent with one half of ping-pong. We do not see transfer in the other direction but our bead-based assay is not well suited to studying this process, as we do not have excess Spir or Capu in solution (The original observation of ping-pong was made with all proteins in solution (Montaville et al., 2014)). Importantly, the intermediate phenotype we observe in flies expressing Spir(A*)-GFP (Fig. 1F **and** Table 1) could indicate that, while not necessary, ping-ponging may enhance mesh assembly in the oocyte. Perhaps Spir helps Capu fall off of filaments when they reach their target. Additionally, Spir could pass the filament to another protein that anchors it at the vesicle even more stably, influencing the dynamics of the mesh. It is also possible that capping serves an important role in vivo, independent of the processes we are studying here.

The classical nucleation promoting factors (NPFs), N-Wasp and Scar, bind actin monomers and the Arp2/3 complex to stimulate nucleation by the Arp2/3 complex (Machesky et al., 1999; Welch and Mullins, 2002). Similarly, the yeast NPF, Bud6, binds an actin monomer and the formin Bni1 to stimulate nucleation by Bni1 (Park et al., 2015; Graziano et al., 2011; Tu et al., 2012). In each of these cases, the NPF has negligible independent activity. In contrast, APC and Spir can nucleate alone, as well as synergize with a formin (Moseley et al., 2007; Okada et al., 2010; Quinlan et al., 2005; Montaville et al., 2014). We note that Spir’s nucleation activity, in the presence of profilin, correlates with synergy: in the absence of nucleation by SpirNT(A*B*C*D*), no synergy is observed; when nucleation is augmented, (SpirNT(A*) nucleation is ∼3x stronger than SpirNT), synergy with Capu is also enhanced over wild type (∼4x). These are only two data points, but they suggest that Spir’s nucleation contributes to synergy, as opposed to Spir acting as a passive actin binding protein, or NPF. In fact, if monomer delivery were Spir’s role, one might expect SpirNT(A*) to be a weaker Capu activator because it has fewer functional WH2 domains. Thus, we conclude that Spir and Capu collaborate by a mechanism similar to that proposed for APC/mDia1 – Spir nucleates seeds that are elongated by Capu (Fig. 7B).

### In vivo implications

The filament orientation we observe, barbed-ends away from the bead, reflects the fact that Spir is attached to the beads and Capu is bound to Spir in our assays. As we and others have shown, when a formin is bound to a bead, filaments grow with their barbed ends at the bead surface. Both mSpire and Fmn-2 are enriched on Rab-11 vesicles. We speculate that Fmn2 is affiliated by binding mSpire as opposed to an independent association with the vesicle. We base this on the finding that mSpire binds directly to membranes and the fact that, in flies, GFP-Capu is diffuse throughout the oocyte (Tittel et al., 2015; Quinlan, 2013). If we are correct, then the geometry in our assays mimics the situation *in vivo* and indicates that the actin filaments are oriented opposite to what was originally proposed by others (Schuh, 2011; Montaville et al., 2014). Long distance transport is still possible with this orientation of filaments (Fig. 7C). Instead of myosin V capturing filaments from a neighboring vesicle, Spir or another protein could capture a growing filament and myosin V on the vesicle from which the filament originated could walk along the filament pulling the vesicle towards its neighbor (Fig. 7C’). In this case, pointed end attachment would not have to be as long lasting as barbed end retention. These conditions are surprisingly well satisfied in our simplified system.

Increased Capu expression is sufficient to build a mesh in the absence of Spir or in the absence of direct interaction with Spir (Dahlgaard et al., 2007; Quinlan, 2013). In these cases, the mesh is not as dense or as long lived. No mesh is built when exogenous Spir expression is driven in the absence of Capu (Dahlgaard et al., 2007). Thus we propose that Spir’s primary role is to enhance Capu’s actin assembly activity in vivo. The fact that Spir(A*)-GFP expression partially rescues loss of Spir indicates that barbed-end binding by Spir is not necessary. We note, however, that actin nucleation is enhanced, which could compensate for loss of ping-ponging or other barbed-end binding roles of Spir. Why do formins need NPFs? In addition to adding another level of control in the cell, nucleation by many formins is dampened by profilin. In the case of the fly oocyte, we also note that the cytoplasm is enriched with microtubules which potently inhibit Capu by binding to its tail, where the Spir-KIND domain also binds (Roth-Johnson et al., 2014). Thus Spir is well suited to simultaneously protect Capu from inhibitory factors and amplify its activity.

## Materials and Methods

### DNA constructs

CapuCT (aa 467-1059) constructs were expressed from a modified pET15b vector with an N-terminal hexahistidine tag (Vizcarra et al., 2011). All other proteins were expressed from a modified pET20b(+) vector with no affinity tag. A native poly-histidine region within the Spir-KIND domain is sufficient for binding of these constructs to TALON® resin (Clontech).

Gibson cloning was employed for the scar-free introduction of Avidity’s 45 bp Avitag™ (translated sequence: GLNDIFEAQKIEWHE) to all proteins requiring biotinylation for bead-conjugation. This tag was introduced to the C-terminus of all constructs.

### Expression, purification, and biotinylation of proteins

Actin was purified from *Acanthamoeba castellani* as described (Zuchero, 2007) and stored in G-buffer (2 mM Tris-Cl, pH 8.0, 0.1 mM CaCl_2_, 0.2 mM ATP, 0.5 mM TCEP, 0.04% sodium azide). Expression was induced in Rosetta™(DE3) cells (EMD Biosciences). Bacteria were grown in Terrific Broth medium supplemented with 100 mg⁄L ampicillin and 32 mg⁄L chloramphenicol. Cells were grown to an OD of 0.6 at 37°C, cooled to 18°C for 1 hour, induced with 250 μM IPTG, and shaken for 18 hours at 18°C. Bacteria were harvested by centrifugation. Pellets were washed once with ice-cold PBS, flash-frozen in liquid nitrogen, and stored at −80°C.

All purification steps were carried out at 4°C or on ice. Thawed cells were diluted at least two-fold with lysis buffer (50 mM sodium phosphate pH 8.0, 1 mM β-ME, 300 mM NaCl) supplemented with 1 mM phenylmethylsulfonyl fluoride (PMSF) and 2 μg⁄mL DNaseI and then lysed by microfluidizing, 2-3x. Cell debris was removed by centrifugation at 20,000×g for 20min at 4°C. Clarified lysates were then rocked with TALON® resin for 1 hour at 4°C (4 mL slurry for every 1 L culture pellet). The TALON® resin was washed with 20 column volumes of lysis buffer, followed by washing with 20 column volumes of wash buffer (lysis buffer, at pH 7.0). Proteins were eluted with elution buffer (wash buffer, plus 200 mM imidazole), until little or no protein remained on the column, as determined by a Coomassie-stained dot-blot of the eluates.

TALON® eluates were pooled and dialyzed 2 x 2 hours and once overnight against 1 L volumes of 10 mM Tris, 1 mM DTT, pH 8.0; or, 5 mL of the most concentrated eluates were buffer exchanged into the same, using a PD-10 desalting column (GE Life Sciences). Protein was loaded onto a MonoQ anion exchange column (GE Life Sciences) and eluted using a gradient of 50–500 mM KCl over 60 column volumes for Spir-KIND, 50–250 mM KCl over 100 column volumes for CapuCT, or 0–500 mM KCl over 60 column volumes for SpirNT. Pooled fractions from the MonoQ column were again dialyzed or buffer exchanged as described above. Unless tagged with an Avitag™, 50% glycerol was added to the overnight dialysis step (1:1 glycerol:buffer). The protein was flash-frozen in liquid nitrogen in 10-50 μL aliquots and stored at −80°C.

If tagged with an Avitag™, 2 mL of the protein was added to 223 μL Biomix B and 5 μL (5 μg) BirA (both from Avidity) and rocked overnight at 4°C. The reaction was again loaded onto a MonoQ anion exchange column and eluted as described above. Pooled fractions from the MonoQ column were dialyzed or buffer exchanged as described above, with 50% glycerol added to the overnight dialysis step and flash-frozen in liquid nitrogen.

Spir-KIND, SpirNT, and CapuCT concentrations were calculated based on their absorbances at 280nm (ε_280_ = 17,452 cm^−1^M^−1^ for KIND, 25,575 cm^−1^M^−1^ for Spir-NT, and 75,200 cm^−1^M^−1^ for Capu-CT) (Quinlan et al., 2007).

Purified VopL and mDia1-CT proteins were generously provided by the Kovar lab (U Chicago).

### Pyrene actin assembly assays

Bulk actin assembly assays were carried out essentially as described (Zuchero, 2007). Briefly, actin (5% pyrene labeled) was incubated for 2 min at 25°C with 200 μM EGTA and 50 μM MgCl_2_ to convert Ca-actin to Mg-actin. When included in the experiment, *Schizosaccharomyces pombe* profilin (typically 2:1 profilin:actin) was incubated with actin for 2 min at 25°C before conversion to Mg-actin. Polymerization was initiated by adding polymerization buffer (KMEH, final concentration: 10 mM HEPES, pH 7.0, 1 mM EGTA, 50 mM KCl, 1 mM MgCl_2_) to the Mg-actin. Additional components, such as CapuCT, SpirNT, and capping protein were combined in the polymerization buffer before addition to Mg-actin. Fluorescence was monitored in a TECAN F200 with λ_ex_ = 360 ± 17 nm and λ_em_ = 400 ± 10 nm.

Actin seeds were prepared by polymerizing 5 μM actin at 25°C for 1 hour in KMEH. The filaments were dispensed in 5 μL aliquots and allowed to reequilibrate for 2–3 hours at 25°C. SpirNT was incubated with filaments for 3 min at 25°C. During this incubation time, monomeric actin was converted to Mg-actin. Using a cut pipette tip to prevent shearing, polymerization buffer was added to Mg-actin and then mixed with seeds plus SpirNT. The slope of the pyrene fluorescence trace between 200 and 500 s was considered the elongation rate.

### TIRF microscopy assays

Coverslips were prepared and functionalized with polyethylene glycol (final surface composition, 97% methoxy-PEG and 3% biotin-PEG; JenKem Technology, Allen, TX) as previously described (Bor et al., 2012). Biotinylated coverslips were stored in a sealed container at 4°C for up to 2 months before use. Flow cells with volumes of ∼10–15 μl were assembled using thin strips of double-stick tape. Streptavidin-coated microspheres were either colorless or Flash Red fluorescent, with mean diameters of ∼100 nm or ∼1 μm (Bangs Laboratories), respectively. Spheres were washed 2x with ∼20 volumes of KMEH, resuspended in 1 volume of KMEH + 1 mg/mL BSA, and added in 10 μL aliquots to 40 μL pre-mixed protein mixtures in KMEH. After a 10 min incubation on ice, spheres were spun for 10 min at 10,000x RPM and 4°C. Pellets were gently resuspended in 30 μL KMEH. Pellets were briefly sonicated (∼5 s) if clumped and not well-dispersed when visualized by TIRF microscopy.

All buffers were flowed into cells in 25 μL volumes in the following order: 1) blocking buffer (1x PBS, pH 8.0, 1% pluronic, 0.1 mg/mL casein) with 2 min incubation; 2) TIRF buffer (50 mM KCl, 1 mM MgCl_2_, 1 mM EGTA, 10 mM HEPES, pH 7.0, 0.2 mM ATP, 50 mM DTT, 0.2% methylcellulose, 20 mM glucose); 3) beads (5 μL resuspended beads, 25 μL TIRF buffer) with 2 min incubation; 4) TIRF buffer; 5) TIRF buffer, supplemented with GCC mix (0.25 mg/mL glucose oxidase, 0.05 mg/mL catalase, 0.8 mg/mL casein), actin (typically 1 μM, 20% Oregon Green labeled), and profilin and/or capping protein, when appropriate. In this final step, actin and profilin (when present) were mixed and incubated for 1 min, then added to the other buffer components immediately before being flowed into the cell.

Time zero was defined as the moment the final 25 uL mix, including actin, was entirely added. Polymerization was visualized immediately (typically, t = ∼30 s) on a DMI6000B TIRF microscope (Leica, Wetzlar, Germany), controlled by LAS X (Leica software). Images were acquired with a DU897 EMCCD camera (Andor) and 100x/1.47 HCX PL APO objective (Leica) at ∼25°C. All analyses were performed on raw data in FIJI. The brightness and contrast of figure images were minimally adjusted for clarity of image features. Filament lengths, elongation rates, and kymographs were analyzed/prepared with JFilament incorporated in FIJI (Smith et al., 2010). Plots were made and statistical analyses conducted in Kaleidagraph. Filament dwell times were obtained by manually tracking individual, retained filaments until clearly released from a bead. Some filaments did not become clearly visible in the TIRF plane until some growth had already occurred. In these cases, time was added to the measured retention period, proportional to the initial length of the filament and the average filament elongation rate observed in the experiment.

For 2-color experiments, 1 μM Cy3B-labeled actin was introduced to the functionalized, bead-bound flow cell. Following this (step 5 of the aforementioned protocol), and after allowing beads to briefly (1–2 mins) polymerize the Cy3B-actin, the following components were flowed in: 6) TIRF buffer supplemented with 1 μM phalloidin, with 2 min incubation; 7) TIRF buffer supplemented with GCC mix, 600 nM 20% labeled OG-actin, and 300 nM profilin. Polymerization was visualized as described above.

For the plots generated in Fig. 6D/E, circles of 10 and 11 px radii (1.6 and 1.76 μm, respectively) were drawn around the centroids of each bead. Actin fluorescence was measured within these circles over time. The integrated density measurements of the 10 px circles were subtracted from those of their concentric, 11 px circles, to measure 1 px-wide (160 nm) bands of actin fluorescence, 1.6 μm away from beads. A script that performs these operations in FIJI is available upon request.

Fluorescence measurements in Fig. 6D/E are expressed as a fraction of their initial values (i.e. a 10% increase = 1.1). Discrepancies in initial fluorescence values for different bead types were observed, in part, due to real differences in assembly rates, because actin polymerization could not be observed instantly. To scale these differences in assembly prior to image collection, the average starting values of each bead type were compared. The larger ratio of these two values was used as a multiplier for the more active bead type. For example, if measurements of SpirNT(A*)- and SpirNT-beads were 110% their initial values, these quantities would each be plotted as “1.1.” However, if the average, initial values of SpirNT(A*)-beads were twice those of SpirNT-beads, the quantities would be plotted as “2.2” and “1.1,” respectively, reflecting the 2-fold greater assembly by SpirNT(A*)-beads prior to image collection.

### Drosophila stocks

The following stocks were obtained from the Bloomington Drosophila Stock Center (NIH P40OD018537): *spir*^1^ and *capu*^1^ (Manseau and Schüpbach, 1989); Df(2L)Exel^6046^ (Exelixis); and nos-GAL4-vp16 (Van Doren et al., 1998). Mutant SpirB-GFP transgenes were generated by inserting the coding region of spir (CG10076-RB) with point mutations created by QuikChange mutagenesis, between the KpnI and SpeI sites of pTIGER (Ferguson et al., 2012) with mEGFP inserted between the BamHI and XbaI sites. pTIGER plasmids were integrated at the attP2 landing site by BestGene.

### Fertility assays

Approximately 100 test females were crossed to 40 wild-type males and kept on apple plates for 2 nights at 25°C. Flies were pre-cleared on a fresh plate with yeast paste for at least 1.5 hours, the plate was changed and eggs laid over the next 3 hours were collected. Typically, 100 eggs were laid in this time period. Eggs were transferred to a fresh plate and stored at 25°C. The number of eggs that hatched after 24 hours was recorded. Each trial was repeated at least three times with independent crosses.

### Fly oocyte microscopy and staining

The visualization of cytoplasmic flows and the actin mesh were performed on a Leica SPE I inverted confocal microscope. Flies were kept at 25°C and fed yeast paste for ∼24 hours before an experiment. Flows were visualized by imaging autofluorescent yolk granules of egg chambers teased apart in Halocarbon oil 700. The actin mesh was stained as described by Dahlgaard et al. (Dahlgaard et al., 2007) with modifications. Briefly, ovaries were dissected, teased apart and fixed in 10% paraformaldehyde/PBS (pH 7.4) for a total of less than 20 minutes. Samples were stained with 1 μM AlexaFluor488-phalloidin diluted in 0.3% Triton X-100/PBS for 25 minutes at room temperature. Samples were then washed extensively and mounted in ProLong Gold with DAPI. Images were recorded within 24 hours of staining because phalloidin staining quality degraded over time, as has been reported (Dahlgaard et al., 2007).

---

**Fig. S1.**
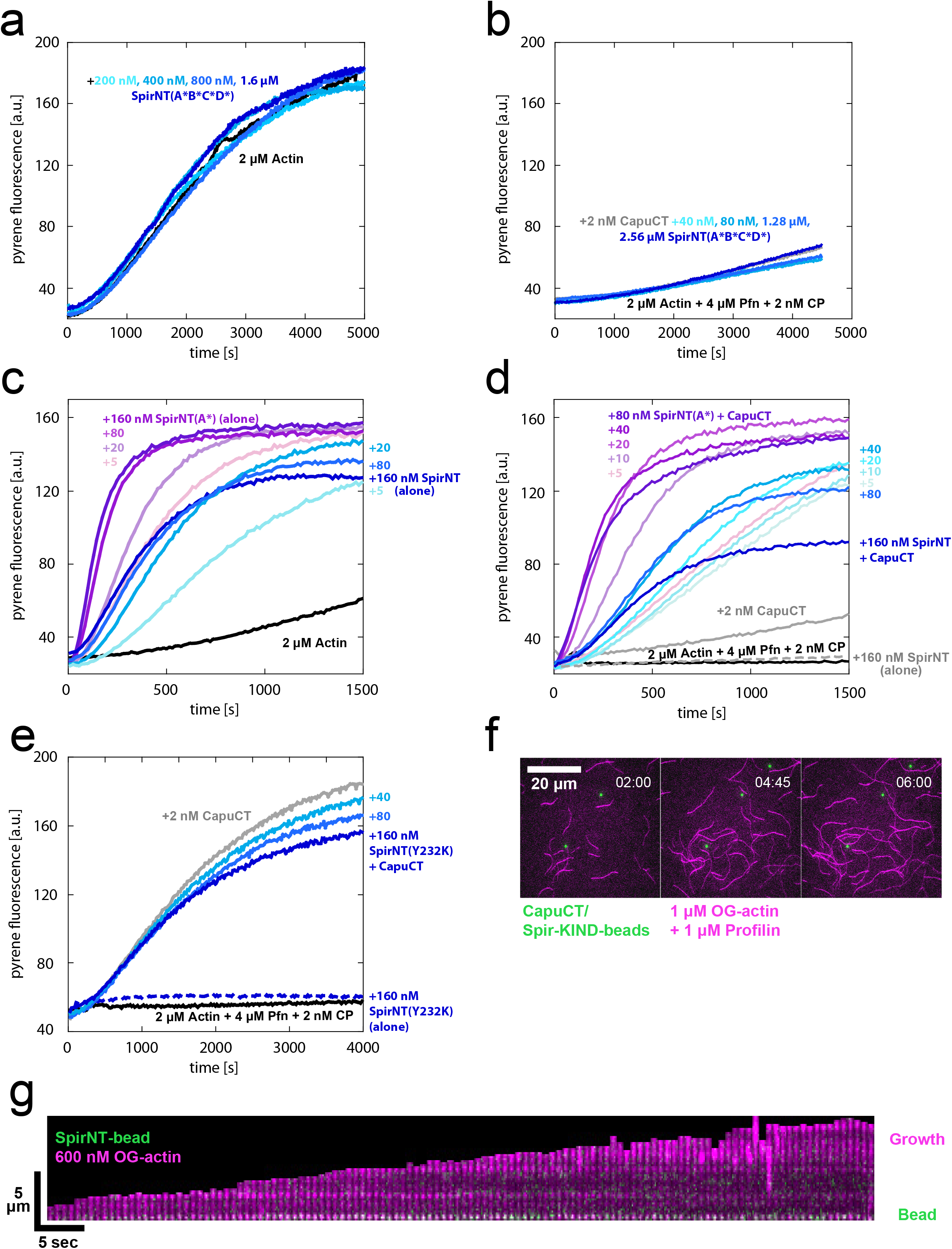
*(A)* SpirNT(A*B*C*D*) does not assemble actin. *(B)* SpirNT(A*B*C*D*) does not synergize with CapuCT to assemble actin. *(C)* SpirNT(A*) is a superior actin nucleator and does not sequester actin like wild-type SpirNT. *(D)* SpirNT(A*) demonstrates enhanced synergy/actin assembly with CapuCT and does not sequester actin like wild-type SpirNT. *(E)* A Capu-binding mutant, SpirNT(Y232K), does not synergize with CapuCT to assemble actin. *(F)* Spir-KIND/CapuCT-beads do not assemble actin or bind actin filaments. *(G)* Retention of nucleated filaments by SpirNT-beads is not dependent on the presence of CapuCT or profilin. Fiducial marks (bright, horizontal lines) in the kymograph are not displaced as the filament elongates, indicating growth away from the bead. See also, *Video 7*.

## Video Legends

**Video 1** – Nucleation by CapuCT/SpirNT-beads

Video corresponds to TIRF images shown in Fig. 2A. 1 μm diameter, Flash Red streptavidin-beads (green channel) were conjugated to SpirNT-biotin, incubated with CapuCT, and allowed to polymerize 1 μM, 20% labeled OG-actin (magenta channel), in the presence of 1 μM *S. pombe* profilin. Actin filaments rapidly emanate from beads.

**Video 2** – *Retention by CapuCT/SpirNT-beads*

TIRF video of 100 nm diameter streptavidin-beads conjugated to SpirNT-biotin, incubated with GFP-CapuCT (green channel), and allowed to polymerize 600 nM, 20% labeled OG-actin (magenta channel). The low concentration of actin inhibits spontaneous nucleation and growth at filament pointed ends. Two beads are each seen to retain several, nucleated filaments.

**Video 3** – *Retention by CapuCT/SpirNT-bead in Presence of Profilin*

Video corresponds to TIRF montage and kymograph (white arrow indicates tracked filament) in Fig. 2B. 100 nm diameter streptavidin-beads were conjugated to SpirNT-biotin, incubated with GFP-CapuCT (green channel), and allowed to polymerize 600 nM, 20% labeled OG-actin (magenta channel), in the presence of 600 nM *S. pombe* profilin. The low concentration of actin and presence of profilin strongly inhibit spontaneous nucleation and growth at filament pointed ends. The central bead is seen to retain several, nucleated filaments.

**Video 4** – *Elongation by mDia-bead*

Video corresponds to TIRF kymograph (white arrow indicates tracked filament) in Fig. 2C. 1 μm diameter, Flash Red streptavidin-beads (green channel) were conjugated to mDia1, and allowed to polymerize 1 μM, 20% labeled OG-actin (magenta channel), in the presence of 1 μM *S. pombe* profilin. A bead nucleates and accelerates the elongation of a filament for several minutes.

**Video 5** – *Nucleation by SpirNT-beads*

Video corresponds to TIRF images shown in Fig. 3A. 1 μm diameter, Flash Red streptavidin-beads (green channel) were conjugated to SpirNT-biotin and allowed to polymerize 1 μM, 20% labeled OG-actin (magenta channel). Actin filaments rapidly emanate from beads.

**Video 6** – *Retention by SpirNT-bead in Presence of Profilin*

Video corresponds to TIRF kymograph (white arrow indicates tracked filament) in Fig. 3C. 1 μm diameter, Flash Red streptavidin-beads (green channel) were conjugated to SpirNT-biotin and allowed to polymerize 1 μM, 20% labeled OG-actin (magenta channel), in the presence of 1 μM *S. pombe* profilin. The low concentration of actin inhibits spontaneous nucleation and growth at filament pointed ends. The bead is seen to briefly retain several nucleated filaments.

**Video 7** – *Retention by SpirNT-bead (Without Profilin or CapuCT)*

Video corresponds to TIRF kymograph (white arrow indicates tracked filament) in *Fig. S1G*. 100 nm diameter streptavidin-beads were conjugated to SpirNT-biotin and mCherry-biotin (green channel), and allowed to polymerize 600 nM, 20% labeled OG-actin (magenta channel). The low concentration of actin inhibits spontaneous nucleation and growth at filament pointed ends. The bead is seen to retain a nucleated filament.

**Video 8** – *Barbed End Capture by SpirNT-bead*

Video corresponds to TIRF montage and kymograph (white arrow indicates tracked filament) in Fig. 4A. 100 nm diameter streptavidin-beads were conjugated to SpirNT-biotin and allowed to polymerize Cy3B-actin (green channel). After briefly incubating the Cy3B-labeled filaments with phalloidin, 600 nM, 20% labeled OG-actin (magenta channel) and 300 nM *S. pombe* profilin were flowed in to replace the Cy3B-actin monomers. The low concentration of actin and the presence of profilin strongly inhibit spontaneous nucleation and growth at filament pointed ends, effectively labeling barbed ends in magenta. A free filament diffuses and grows until it is captured by a SpirNT-bead (open triangle). The filament resumes growth once released by the bead (solid triangle).

**Video 9** – *Retention and Capture by CapuCT/SpirNT-beads*

Video corresponds to TIRF montage in Fig. 4B. 100 nm diameter streptavidin-beads were conjugated to SpirNT-biotin, incubated with GFP-CapuCT (green channel), and allowed to polymerize 600 nM, 20% labeled OG-actin (magenta channel), in the presence of 2.4 μM *S. pombe* profilin and 600 pM mouse capping protein. The low concentration of actin and the presence of profilin strongly inhibit spontaneous nucleation and growth at filament pointed ends. The presence of capping protein inhibits growth at barbed ends that are unbound by CapuCT. Two nucleated filaments (cyan triangles) are retained by their nucleating bead (bottom-right) for minutes, while their CapuCT-bound, fast-growing barbed ends are captured by other CapuCT/SpirNT-beads (gold triangles; top-center and top-left).

**Video 10** – *Nucleation by SpirNT-beads in Presence of Profilin (WT vs. A*)*

Video corresponds to TIRF images shown in the Fig. 5D montages. 1 μm diameter, Flash Red streptavidin-beads (green channel) were conjugated to SpirNT-biotin or SpirNT(A*)-biotin and allowed to polymerize 1 μM, 20% labeled OG-actin (magenta channel), in the presence of 1 μM *S. pombe* profilin. The SpirNT(A*)-bead (right) is seen to nucleate and retain many more filaments than the SpirNT-bead (left).

## Acknowledgements

We acknowledge all members of the Quinlan lab for their tireless support and thank David Kovar for generously gifting us purified, SNAP-tagged mDia1.

For the duration of this work, A.O. Bradley was funded by NIH-supported CMB Training Grant at UCLA. M.E. Quinlan is supported by ##.

The authors declare no competing financial interests.

## Author Contributions

Margot E. Quinlan (lead) and Christina L. Vizcarra (supporting) conceptualized the experiments. Alexander O. Bradley (lead) and C.L. Vizcarra (supporting) developed the methodology. A.O. Bradley performed and analyzed all experiments; with the exception of the genetics experiments presented in Table 1, performed by Hannah M. Bailey. A.O. Bradley prepared all figures. M.E. Quinlan and A.O. Bradley wrote the manuscript. All authors contributed to discussion and evaluation of the manuscript.

